# Prelimbic cortex perineuronal net expression and social behavior: Impact of adolescent intermittent ethanol exposure

**DOI:** 10.1101/2024.03.11.584421

**Authors:** Trevor T. Towner, Harper J. Coleman, Matthew A. Goyden, Andrew S. Vore, Kimberly M. Papastrat, Elena I. Varlinskaya, David F. Werner

## Abstract

Adolescent intermittent ethanol (AIE) exposure in rats leads to social deficits. Parvalbumin (PV) expressing interneurons in the prelimbic cortex (PrL) contribute to social behavior, and perineuronal nets (PNNs) within the PrL preferentially encompass PV interneurons, acting as significant regulators of these cells. AIE exposure increases PNNs, however, it is unknown if this upregulation contributes to AIE-induced social impairments. The current study was designed to determine the effect of AIE exposure on PNN expression in the PrL and to assess whether PNN dysregulation contributes to social deficits elicited by AIE. cFos-LacZ male and female rats were exposed every other day to tap water or ethanol (4g/kg, 25% w/v) via intragastric gavage between postnatal day (P) 25-45. We evaluated neuronal activation by β-galactosidase expression and PNN levels either at the end of the exposure regimen on P45 and/or in adulthood on P70. In addition, we used Chondroitinase ABC (ChABC) to deplete PNNs following adolescent exposure (P48) and allowed for PNN restoration before social testing in adulthhod. AIE exposure increased PNN expressionin the PrL of adult males, whereas a decrease in PNNs was evident immediately following AIE. Vesicular glutamate transporter 2 (vGlut2) and vesicular GABA transporter (vGat) near PNNs were moderately downregulated only in AIE-exposed females. Gene expression of PNN components were largely unaffected by AIE exposure. Removal and reestablishment of PNNs in the PrL led to upregulation of PNNs and social impairments in males, regardless of adolescent exposure. These data suggest that AIE exposure in males produces an upregulation of PrL PNNs that likely contribute to social impairments induced by AIE.

## Introduction

Adolescence is the critical developmental period following childhood during which substantial maturation of the body and brain occurs (Spear, 2000), ultimately preparing an individual for adulthood. The adolescent brain undergoes significant experience-driven synaptic plasticity (Selemon, 2013), refinement of critical circuitry, such as dopaminergic innervation of the prefrontal cortex (Wahlstrom et al., 2010), and pruning of excess synapses (Rakic et al., 1994; Spear, 2013). Adolescents commonly display increased impulsivity and risk-taking (Romer, 2010), two behaviors that likely contribute to the high prevalence of initiating alcohol use during this developmental period (Hamilton et al., 2014).

Adolescents typically drink in more risky patterns such as binge (4-5 drinks per occasion for females and males, respectively) and high intensity drinking (10+ drinks per occasion) (Patrick & Terry-McElrath, 2019). Between the ages of 12 and 25, binge drinking rates drastically increase, with nearly 30% of individuals reporting at least one instance of bingeing in the previous month (SAMHSA, 2022). Importantly, two strong predictors of developing alcohol use disorder (AUD) are early initiation of alcohol use (Hingson et al., 2006) and elevated levels of heavy alcohol consumption (Nieto et al., 2021). Thus, adolescents who initiate alcohol use at an early age and drink in high quantities are uniquely vulnerable to the later development of problematic drinking and/or AUD.

Short-term adverse consequences of binge drinking are noted in cognitive performance (Hanson et al., 2011; Mahedy et al., 2018) and may be associated with altered gray and white matter maturation (Lees et al., 2020). However, our understanding of the long-term consequences of adolescent binge drinking in humans is less clear, with some evidence of prolonged cognitive deficits after alcohol cessation (Carbia et al., 2017). Alternatively, the behavioral consequences of adolescent intermittent ethanol (AIE) exposure of laboratory rodents have been extensively studied in recent years (Crews et al., 2019).

Our lab has repeatedly found that AIE exposure induces social deficits specifically in male rats, with no changes evident in females (Dannenhoffer et al., 2018; Towner et al., 2022a; Varlinskaya et al., 2020; Varlinskaya et al., 2014). We recently identified socially evoked neuronal activity within the prelimbic cortex (PrL) was lower in AIE-exposed males compared to water-exposed controls (Towner et al., 2023). To determine whether this altered activation state was functionally related to AIE-induced social deficits, we used a pharmacogenetic approach that allows inactivation of previously activated PrL neuronal ensembles with Daun02 in cFos-LacZ rats. Interestingly, inactivation of the PrL reduced social interaction in controls, but failed to alter social behavior of AIE-exposed males, suggesting a potential dysfunction in recruitment of PrL neurons by social interaction.

The PrL is particularly vulnerable to developmental ethanol exposure due to substantial remodeling during adolescence (Larsen & Luna, 2018). For example, AIE-exposed subjects display reduced dopaminergic (Boutros et al., 2015; Obray et al., 2022; Shnitko et al., 2016; Trantham-Davidson et al., 2017), cholinergic (Fernandez & Savage, 2017), and astrocytic (Healey et al., 2022) function in the PrL. Notably, AIE also impairs inhibitory signaling in the PrL as illustrated by reduced gamma aminobutyric acid (GABA) receptor function (Centanni et al., 2017) and decreased fast-spiking interneuron (FSI) intrinsic excitability (Trantham-Davidson et al., 2017). These AIE-associated alterations of GABAergic tone may contribute to differential neuronal activation of the PrL after an interaction with a social partner (Towner et al., 2022a).

Inhibitory interneurons control the excitatory tone by synapsing on pyramidal cells (Blatow et al., 2003; Hartwich et al., 2009), and in vivo electrophysiology has identified an importance of GABAergic signaling during social interaction (Filiano et al., 2016; Liu et al., 2020; Scheggia et al., 2020; Xu et al., 2019; Zhao et al., 2022). For example, roughly 75% of the PrL parvalbumin (PV) expressing FSIs had increased firing rates when the experimental subject was interacting with a social stimulus (Liu et al., 2020).

Perineuronal nets (PNNs) are part of the extracellular matrix (ECM) and preferentially encompass PV expressing FSIs in the cortex (Carceller et al., 2020; Fawcett et al., 2019). Chondroitin sulfate proteoglycans (CSPGs) such as aggrecan, brevican, neurocan, and phosphacan, are the primary components of PNNs and aid in the lattice-like formation of these structures (Dyck & Karimi-Abdolrezaee, 2015; Rowlands et al., 2018). PNNs are developmentally regulated, with rapid increases in PNN expression detected during the juvenile to adolescent transition and extending into adulthood (Baker et al., 2017; Drzewiecki et al., 2020; Mauney et al., 2013; Ueno et al., 2018). Thus, it is not surprising that PNN expression is altered by AIE, with findings of upregulated CSPGs in the orbitofrontal cortex (OFC) (Coleman et al., 2014). This upregulation of PNNs in the OFC was recently replicated and extended to the medial prefrontal cortex, with a similar increase evident after AIE (Dannenhoffer et al., 2022). However, the studies that have identified AIE-induced alterations of PNNs included only male subjects, with the effects of AIE on PNN expression in females still unknown. In addition, the role of PrL PNN upregulation in contributing to AIE-induced behavioral alterations has not been evaluated.

The present study was designed to assess the impact of AIE exposure on PNN expression in the PrL of male and female rats and evaluate whether PNNs contribute to AIE-induced social deficits. We hypothesized that AIE would increase PNN expression in the PrL regardless of sex, with only AIE-exposed males displaying greater social stimulus-induced activation of PNN-encapsulated cells, supportive of their contribution to AIE-induced social deficits. We further hypothesized that PrL PNN ablation would restore social behavior in AIE-exposed males.

## Materials and Methods

### General Methods

#### Subjects

In all experiments, we used cFos-LacZ transgenic rats that express the E. coli derived LacZ gene driven by a cFos promoter. LacZ encodes β-galactosidase (β-gal) which is a proxy for cFos expression in these animals. The use of these animals as experimental subjects allowed us to measure β-gal expression induced by an interaction with a social partner as a marker for neuronal activation indexed via cFos, similar to what we have done previously (Towner et al., 2022a; Towner et al., 2023). All animals were bred and reared at Binghamton University in temperature-(20-22 °C) and humidity-controlled vivaria with lights on/off at 0700/1900 hours. Water and food were available *ad libitum*. Litters were culled on postnatal day (P) 1 to 8-10 pups, with an equal ratio of males to females whenever possible. On P21, rats were weaned into groups of 3-6 same-sex littermates and ear punched for genotyping, with tissue processing by an outside vendor (TransnetYX). Prior to P25, number of pups per cage was reduced to 3-4, with a variable ratio of +/− genotype for the LacZ gene. Only LacZ+ rats were used for the experiments in the current study, except for Experiment 3 in which LacZ-litter and cage mates were used. Animal housing, care, and experimentation were in accordance with the National Institutes of Health guide for the care and use of laboratory animals and using protocols approved by Binghamton University Institutional Animal Care and Use Committee.

#### Adolescent Intermittent Ethanol Exposure

Starting on P25, rats were exposed every other day to ethanol (25% w/v, 4.0 g/kg) or an isovolumetric equivalent of tap water via intragastric gavage for a total of 11 exposures ending on P45 (see Fig. 1A). We have previously found this ethanol dosing regimen elicits blood ethanol concentrations between 200 mg/dL at the start and ∼125 mg/dL after the final exposure (Kim et al., 2019). All subjects in a cage were assigned to the same exposure condition.

**Figure 1.**
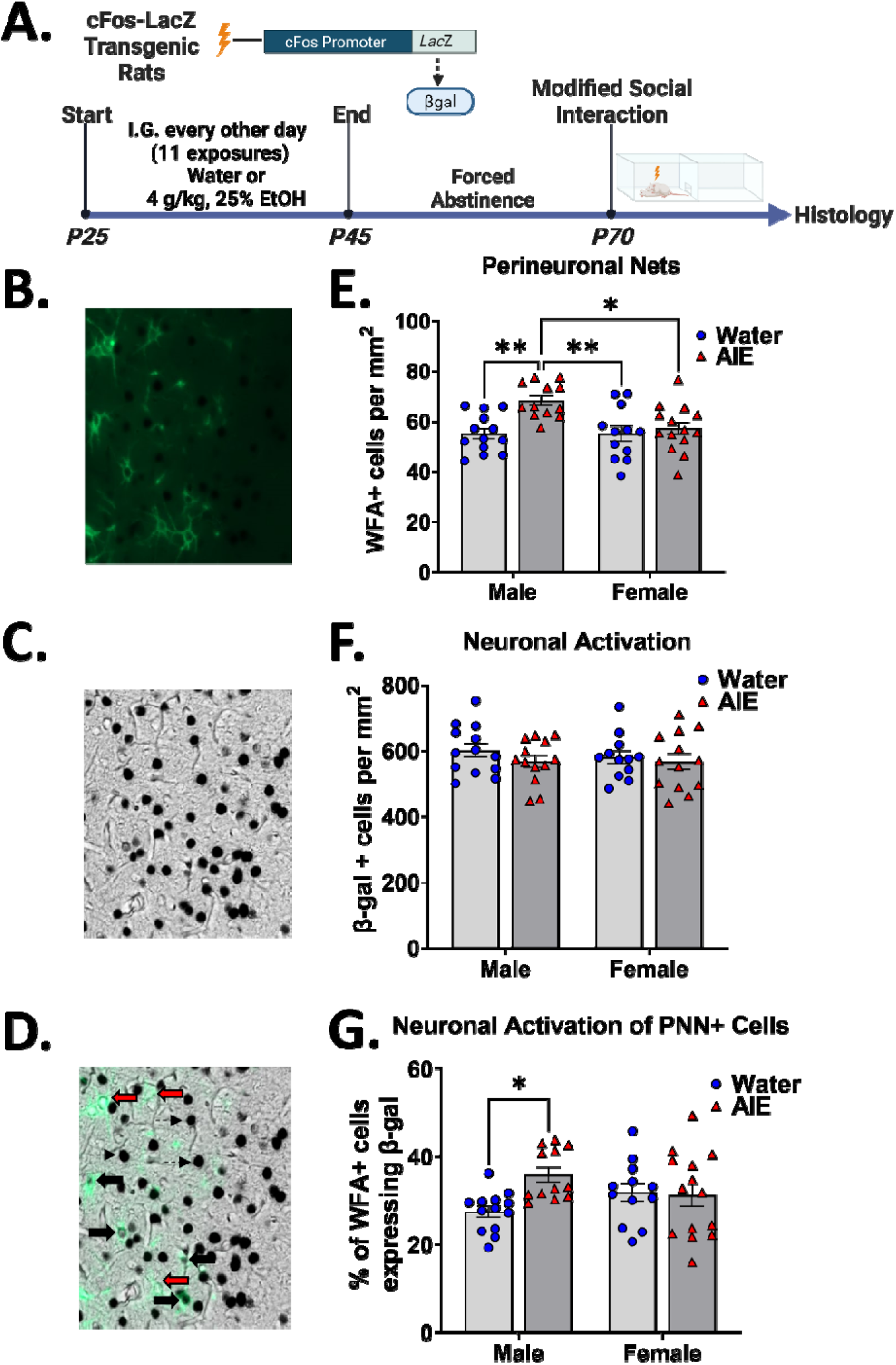
Expression of PNNs and colocalization with neuronal activation following AIE in cFos-LacZ transgenic rats. **A.** Timeline of adolescent water or ethanol (AIE) exposure and social testing in adulthood. **B-D.** Representative images of WFA-labeled PNNs, β-gal, and co-labeling of WFA and β-gal, respectively. **E.** AIE exposure led to an increase in WFA+ cells in the PrL compared to water-exposed males and females of both exposure groups (marked with *). **F.** No differences were found for the number of β-gal+ cells in the PrL of socially tested rats. **G.** A larger percentage of WFA+ cells were activated in AIE-exposed males relative to water-exposed controls. * *p* < 0.05 and ** *p* < 0.01.

#### Modified Social Interaction Test

Social behavior was assessed using a modified social interaction test in an apparatus (46.0 × 30.3 × 30.5 cm) consisting of two equal size compartments separated by a wall with an aperture that allowed for movement between the compartments. Fresh pine shavings were placed at the bottom of the apparatus, and 6% hydrogen peroxide was used to clean each chamber between test sessions. Testing took place under dim lighting (10-15 lux) and in the presence of white noise. Experimental subjects were weighed, marked for identification, and placed in the apparatus for a 30-minute habituation. Following the habituation, a same-sex, unfamiliar adult partner was introduced to the chamber. Social interaction was video recorded for 10 minutes; however, the experimental subject and partner were left together for a total of 60 minutes. This timing was determined in our lab to elicit peak β-gal expression following a stimulus (unpublished parametric findings).

The 10-minute social interaction test was scored by a trained researcher who was blind to the experimental condition of the subjects. Two social behaviors were measured: social investigation and social preference/avoidance coefficient. Social investigation was any instance of sniffing of the social partner. Social preference/avoidance was measured by a coefficient using the following formula: (number of crossovers toward the partner – number of crossovers away from the partner) / (total number of crossovers to and away from the partner) × 100. Positive values of the coefficient are characteristic for social preference, whereas animals with social avoidance have negative values of the coefficient.

#### Statistical Analysis

All dependent variables were assessed using analyses of variance (ANOVAs) in Prism (GraphPad, La Jolla, CA, USA) and represented as means ± standard error of the mean (SEM). In the case of significant main effects and/or interactions, follow up comparisons were conducted with Bonferroni tests. Outliers that were ± two standard deviations outside the mean were removed from analyses.

### Experimental Designs and Procedures

#### Experiment 1. Effects of AIE on PNN expression on activated and non-activated neurons in the prelimbic cortex of socially tested adult rats

To determine whether AIE leads to an upregulation of PNN expression in the PrL, we conducted histology on tissue generated in a previous study (Towner et al., 2023). This experiment used a 2 adolescent exposure (water, AIE) x 2 sex (male, female) design with 10 – 14 LacZ+ animals per group. All animals were exposed to water or AIE as described above and aged into adulthood before social testing to elicit β-gal expression in the PrL (Fig. 1A). Within this study, males exposed to AIE displayed social deficits similar to what we have shown previously (Towner et al., 2022a; Varlinskaya et al., 2020; Varlinskaya et al., 2014), however given our primary interest was to assess PNN expression, and that social response is reported elsewhere, the behavior for these animals is not included in the current study.

##### Tissue Preparation

Following the social interaction, test subjects were intraperitoneally injected with a fatal dose of sodium pentobarbital. Once fully anaesthetized, rats were transcardially perfused with 0.01M phosphate buffered saline (PBS) followed by 4% paraformaldehyde (PFA) in 0.1M phosphate buffer. Brains were extracted and post-fixed in 4% PFA solution for a total of 90 minutes then transferred to a 30% sucrose solution. Once saturated, brains were flash frozen in methyl butane and stored at −80 °C until slicing. Using a Leica CM1860 (Leica Biosystems, Wetzlar, Germany) cryostat, 30μm slices were collected and placed in an antifreeze solution until further processing.

##### X-gal and WFA staining

Every sixth slice containing the PrL (4.40 – 2.52 mm anterior to Bregma) was used and subjected to X-gal staining to label β-gal as we had done previously (Towner et al., 2022a). Briefly, slices were fixed in a in 0.1M PB with 5mM EGTA for 15 minutes and were then washed (twice) for 5 minutes in 0.1M PB. Slices were incubated overnight at 37° C in staining buffer (0.1M PB with 2mM MGCl_2_, 5mM potassium ferrocyanide, and 5mM potassium ferricyanide) and added x-gal stock (5-bromo-4-chloro-3-indolyl-β-D-galactoside in dimethylformamide, 1:50). These same slices were then labeled to identify PNNs. Slices were washed 3 times in PBS for 5 minutes each. Sections were blocked using a carbo-free blocking solution (Vector Laboratories, Burlingame, CA, USA) for one hour at room temperature. Slices were then washed overnight at 4 °C in a biotinylated *Wisteria Floribunda* Agglutinin (WFA, 1:2000, Vector Laboratories, B-1355-2) followed by washing 3x in PBS for 5 minutes before 60-minute incubation with a Streptavidin-conjugated Dylight 488 secondary (1:200, Vector Laboratories, SA-5488). After secondary, slices were again washed in PBS before mounting on charged slides and cover-slipped using fluoromount (Electron Microscopy Sciences, Hatfield, PA, USA).

##### Image Acquisition and Analysis

Slides were imaged using an Olympus VS200 Slide Scanner (Olympus, Center Valley, PA, USA). To visualize WFA-labeled PNNs, images were tiled at 10x using identical exposure timing with a 488-filter cube. These same slices were then imaged with a monochromatic camera under dark-field conditions for assessing β-gal labeled cells. The different channels were separated for analysis of each marker individually and then combined to detect co-labeled cells. This method for labeling β-gal+ cells instead of immunofluorescence was used to ensure similarity in methods between the current and our previous publication (Towner et al., 2022a).

Images containing WFA-labeled PNNs alone were analyzed using Pipsqueak Pro (Rewire Neuro, Portland, OR, USA) with a stock model designed for detection of WFA+ cells. β-gal expression was analyzed using Halo (Indica Labs, Albuquerque, NM, USA) with an algorithm detecting individual somatic markers based on contrast between background and β-gal signal as well as size. For analysis of the co-label of WFA and β-gal, we used the program OlyVIA (Olympus). A 1000 × 400 μm box was cropped from the PrL, spanning all layers of the cortex, and used as the image for the analyses described above. Individual representations of WFA-labeled cells, dark field β-gal labeling, and the co-labeling of WFA and β-gal can be seen in Figure 1B-D.

#### Experiment 2: Long-lasting effects of AIE on PNN-associated excitatory and inhibitory synapses in adulthood

Given that PNNs regulate synaptic plasticity through limiting the removal/addition of new synapses (Bosiacki et al., 2019), we were interested in assessing AIE-induced changes in excitatory and inhibitory synapses in contact with PNNs. The design of this experiment was a 2 adolescent exposure (water, AIE) x 2 sex (male, female) factorial, with n = 10-12 LacZ+ rats per group. Following exposure in adolescence, adult rats were taken directly from the home cage (Fig. 2A) and transcardially perfused with 4% PFA prior to a 90-minute postfix in 4% PFA. Following submersion in 30% sucrose solution, brains were frozen, sectioned (30μm), and stored in antifreeze.

**Figure 2.**
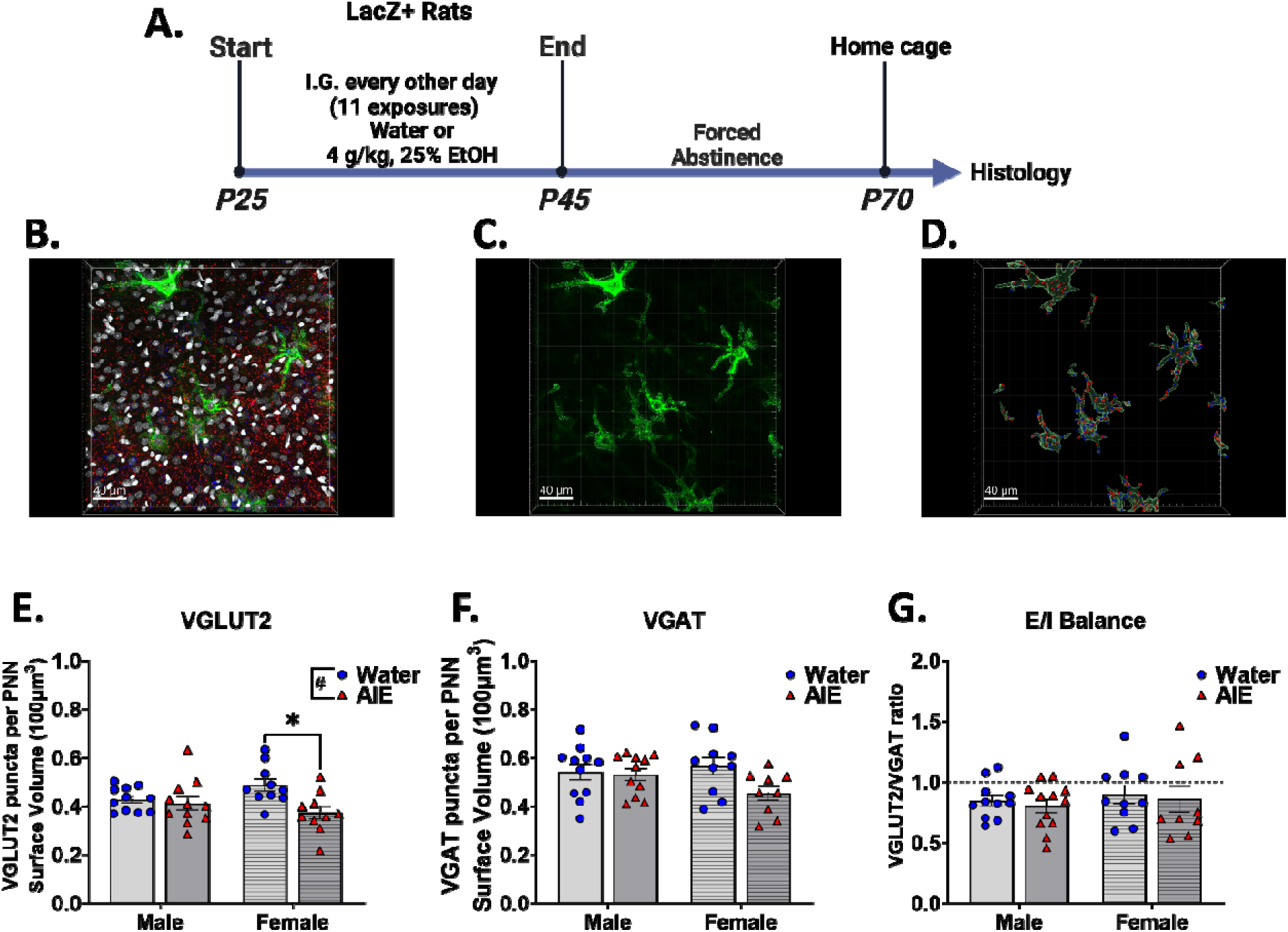
Changes to excitatory and inhibitory synaptic puncta on PNN expressing cells. **A.** Timeline of adolescent exposure and tissue collection for histology. **B-D.** Representative images of labeling PNNs and synaptic puncta (dapi (grey), WFA (green), vGlut2 (red), and vGat (blue)), PNN volume alone, and rendering of synaptic puncta within 1 μm of the PNNs, respectively. **E.** Vglut2 expressing synapses were decreased in AIE-exposed females relative to water-exposed females. **F-G.** No differences in Vgat labeling or excitatory/inhibitory ratio. * *p* < 0.05 indicates difference between water- and AIE-exposed females. # *p* < 0.05 indicates main effect of exposure group. Scale bar represents 40 μm.

##### Immunofluorescent assessments of vesicular glutamate transporter 2 (vGlut2) and vesicular GABA transporter (vGat) containing synaptic puncta

Sections containing the PrL were washed (3x, 5 minutes each), then incubated in carbo-free blocking solution for 60 minutes prior to incubating overnight in WFA (1:2000). After that, sections were washed, blocked with 5% normal donkey serum, 0.2% Triton X-100, 1% Bovine Serum Albumin, and glycine in tris buffered saline (TBS) for 60 minutes. Slices were then transferred to primary cocktail solution containing rabbit anti-vGat (1:2000, Millipore Sigma, AB5062P) and guinea pig anti-vGlut2 (1:2000, Millipore Sigma, AB2251-I) overnight at 4 °C. Then, slices were washed, and incubated in a secondary cocktail solution containing donkey Alexa Flour 680 anti-rabbit (1:1000, Invitrogen, Waltham, MA, USA) and donkey Alexa-Flour 594 anti-guinea pig (1:1000, Jackson Immunoresearch, West Grove, PA, USA) for 60 minutes, then washed again before mounting and cover-slipping with Fluoromount.

Slides were viewed under a Leica TCS SP8 confocal microscope with a 40x objective and images were acquired using LASX navigation software. A single Z-stacked (.35 μm steps) 40x frame (290 × 290 μm) placed in the PrL was captured across 3 sections per subject. Entire Z-stack files were transferred to Imaris (Oxford Instruments, Abingdon, United Kingdom) where images were 3D rendered. PNNs were identified using the surface detection function and traced. Individual vGat and vGlut2 signal within the image were detected using the Imaris spots function. Identified spots that were within 1 μm from the PNNs represented synaptic puncta potentially in contact with the PNN encompassed cell (see Fig. 2B-D). A ratio of puncta per volume of PNNs (100μm^3^) identified was used to normalize across images.

#### Experiment 3: Immediate effects of AIE on WFA-labeled PNNs

This experiment was designed to test whether AIE-induced changes in PNNs could be evident immediately following the AIE exposure using WFA labeling of PNNs in the PrL of animals euthanized following the final gavage (P45, see Fig. 3A). We implemented a 2 adolescent exposure (water, AIE) x 2 sex (male, female) design, with n = 11-12 per group. Animals received their final exposure to either water or ethanol on P45 and were euthanized by rapid decapitation 60 minutes after, with whole brains extracted and postfixed in 4% PFA for 90 minutes. Brains were transferred to 30% sucrose, stored, then processed in accordance with methods described above. Sections containing the PrL were labeled with WFA by washing 3x for 5 minutes each in PBS, incubating for 60 minutes in carbo-free block, incubating overnight at 4 °C in WFA (1:2000), washed again, incubated in secondary (1:200 Streptavidin conjugated Dylight 488), and washed prior to mounting and coverslipping with Fluoromount. Sections were imaged at 10x on an Olympus slide scanner and WFA-labeled cells were detected using Pipsqueak Pro as in Experiment 1.

**Figure 3.**
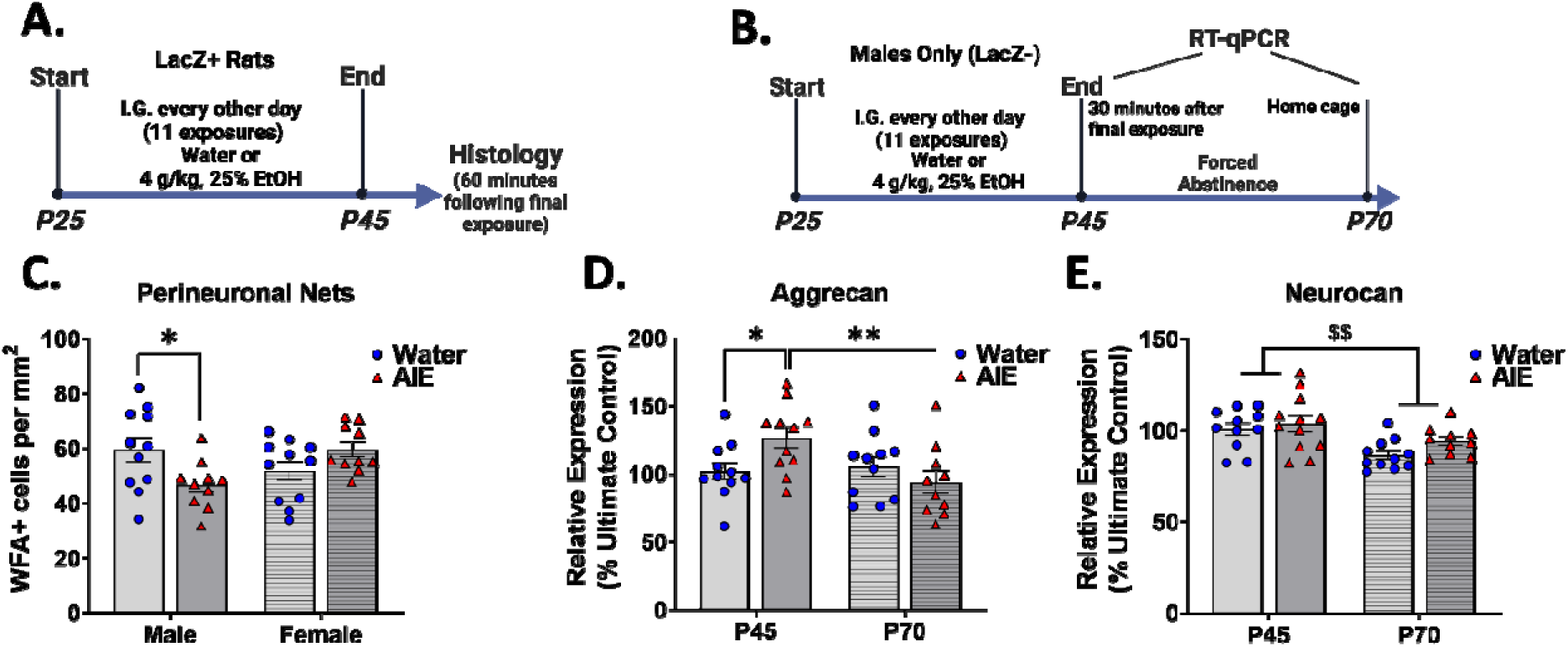
Immediate and long-term effects of AIE on PNN expression in the PrL. **A.** Male and female rats exposed to water or AIE then euthanized for WFA labeling PNNs following the final exposure on P45. **B.** Subjects were exposed to water or AIE and brains extracted for gene expression analysis either 30 minutes following the final exposure on P45 or in adulthood on P70. **C.** AIE-exposed males had lower WFA labeling than water-exposed controls at the end of the exposure regimen. **D.** Aggrecan gene expression was elevated in AIE-exposed males on P45 (immediately following the end of AIE) but not after 25 days of abstinence. **E.** Neurocan gene expression was significantly lower in adults, regardless of adolescent exposure. * *p* < 0.05 and ** *p* < 0.01 indicate difference from AIE-exposed P45 group. $$ *p* < 0.01 indicates difference between ages.

#### Experiment 4: Immediate and long-lasting effects of AIE on PNN-associated chondroitin sulfate proteoglycan (CSPG) gene expression

This experiment tested the immediate and long-lasting effects of AIE on PNNs by measuring expression of chondroitin sulfate proteoglycans genes in the medial prefrontal cortex (mPFC) either during adolescence following the final exposure or in adulthood after 25 days of abstinence. The design was a 2 adolescent exposure (water, AIE) x 2 age (adolescent, adult) factorial, with 9 – 12 male subjects per group. Male rats (LacZ-) were exposed to water or AIE, with brain tissue collected either 30 minutes following the last gavage on P45 or on P70 after 25 days of abstinence (Fig. 3B). Rats were euthanized by rapid decapitation, brains extracted, and immediately flash frozen in methyl butane before storing at −80 °C.

##### Reverse Transcription Polymerase Chain Reaction

Tissue micropunches containing the mPFC (PrL and Infralimbic (IL)) were taken and processed using protocols previously described (Towner et al., 2022b). In short, we used a phenol-chloroform extraction with Qiagen RNAeasy kits (Qiagen, Hilden, Germany) according to the manufacturer instructions. Total RNA was quantified using a spectrophotometer (NanoDrop 2000, Thermo Fisher Scientific, Waltham, MA, USA). Samples were normalized and underwent cDNA synthesis using QuantiTect Reverse Transcription Kits (Qiagen). qPCR was conducted using SYBR green supermix (Bio-Rad, Hercules, CA, USA) to detect amplification in a CFX384 (Bio-Rad) PCR detection system. Relative gene expression was calculated using the delta-delta method (Livak & Schmittgen, 2001) with P45 water-exposed subjects as the ultimate control group. Cyclophilin A gene expression was used as a stable reference gene. The CSPG genes of interest (GOIs) assessed were Aggrecan (Acan), Brevican (Bcan), Neurocan (Ncan), Phosphacan (Pcan), and Versican (Vcan). In addition, we evaluated gene expression changes to metallopeptidases ADAMTs4, ADAMTs5, and MMP9. All forward and reverse primers can be seen in Table 1.

**Table 1.**
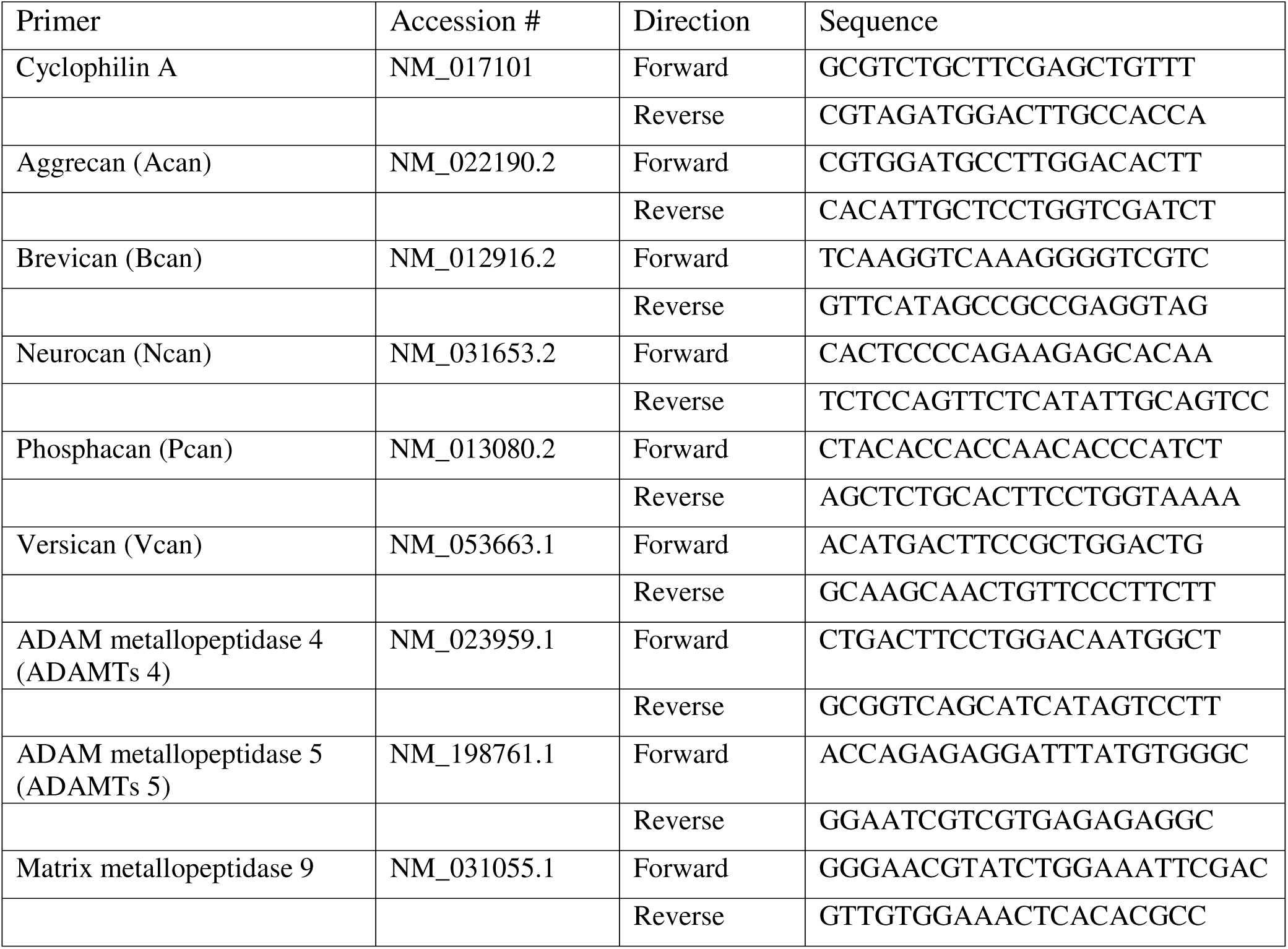
Primer sequences for genes of interest.

#### Experiment 5: Validating ChABC-induced PNN depletion and timing PNN regrowth

The functional role of PNNs in behavior has been established through depletion of PNNs using the enzyme Chondroitinase ABC (ChABC) (Kwok et al., 2008). However, ChABC-induced degradation of PNNs is temporary, and PNNs reform given sufficient time following depletion (Bruckner et al., 1998; Lensjø et al., 2017). In this experiment we wanted to verify that ChABC successfully depletes PNNs and determine the time course of PNN development post-ChABC injection. We assessed PNN levels at three different time points post-ChABC microinjection into the PrL: 7, 14, and 21 days. Both male and females were used, although sex differences in ChABC efficacy have not been reported therefore group sizes were powered based on combining sexes (n = 9-11 per group). Rats were euthanized by rapid decapitation, brains extracted and post-fixed in 4% PFA overnight then processed into 30 μm slices for WFA labeling of PNNs across the PrL. Sections were imaged at 10x using an Olympus slide scanner and WFA+ cells were identified using Pipsqueak Pro.

##### Cannulation and ChABC Injection

Adult rats were anaesthetized with isoflurane (3%) and placed into a stereotaxic frame. During the surgery animals were maintained on 1-3% isoflurane. To target the PrL, bilateral holes approximating the coordinates of AP: +3.2mm, ML: ±0.75mm, and DV: −2.5mm, were drilled into the skull. Bilateral, 26-gauge, steel-guide cannula (PlasticsOne, Roanoke, VA, USA) were placed into the holes. After placing the cannula, animals received either a microinjection of vehicle (PBS) or ChABC (0.09U/μL/side, Millipore Sigma, C3667) with a flow rate of 0.2μL per minute. Injectors were left in place for two minutes to allow diffusion and limit the backflow of drug. Cannulas were then adhered in place by dental cement. Dummy cannulas were inserted and dust caps affixed to the top of the cannula. Rats recovered in cages on top of a warming pad and were given access to food and water. Buprenorphine HCl (0.03mg/kg, intraperitoneal) was administered prior to surgery and every 12 hours following, for a total of 48 hours. Rats were isolate housed for the remainder of the experiment.

#### Experiment 6: Effects of PNN ablation in the PrL on social interaction in adult AIE-exposed rats

We hypothesized that AIE would lead to an upregulation of PNNs as others have shown previously (Coleman et al., 2014; Dannenhoffer et al., 2022) and that this would contribute to aberrant signal processing in the PrL of AIE-exposed subjects that display social impairments. To test whether the alterations in PNN expression contribute to AIE-induced social deficits, we temporarily degraded PNNs in the PrL with ChABC so as to delay the subsequent increases in adulthood following AIE. If AIE increases PNN expression, and we deplete PNNs but permit regrowth in the absence of ethanol, it is possible to restore normal PNN expression as well as function potentially reversing AIE-induced social deficits. To test this prediction, we used a 2 adolescent exposure (water, AIE) x 2 sex (male, female) x 2 drug (vehicle, ChABC) factorial design, with N = 10 – 13 LacZ+ rats per group. Subjects were exposed to water or AIE and three days following the last exposure (P48) underwent cannulation (see cannulation section) of the PrL for PNN depletion. Following three days of isolated recovery, animals were reintroduced to pair-housing with littermates. Approximately 28 days after surgery (P75-78), rats were socially tested and euthanized via transcardial perfusion with brains processed for WFA and β-gal expression (see Experiment 1 x-gal and WFA staining section for method and see Fig. 6A for timeline).

## Results

### Experiment 1. Effects of AIE on PNN expression on activated and non-activated neurons in the prelimbic cortex of socially tested adult rats

To determine whether AIE leads to an upregulation of PNN expression in the PrL, we conducted dual-labeling of β-gal and WFA (Fig. 1B-D). Brain tissue was collected from adult males and female rats exposed to either water or AIE, with all animals exposed to social partners for 60 minutes, with AIE-exposed males demonstrating social deficits indexed via significant decreases in social investigation (see: Towner et al., 2023). A 2 (adolescent exposure) X 2 (sex) ANOVA of the number of WFA+ cells revealed a significant adolescent exposure by sex interaction, F (1,47) = 5.01, *p* < 0.05. As seen in Figure 1E, males exposed to AIE had significantly greater PNN expression than all other groups (*ps* < 0.05), with no other significant differences evident. We also analyzed β-gal expression in the PrL and found that the number of β-gal+ cells did not differ as a function of either adolescent exposure or sex (Fs < 1.24, *ps* > 0.05, Fig. 1F). We also assessed the co-labeling of WFA and β-gal, indicative of activation of PNN encapsulated cells, and found a significant interaction of adolescent exposure and sex, F (1,47) = 5.00, *p* < 0.05, with AIE-exposed males having a significantly greater percentage of WFA+ cells containing β-gal than water-exposed males (Fig. 1G). Post-hoc tests revealed no other group differences.

### Experiment 2: Long-lasting effects of AIE on PNN-associated excitatory and inhibitory synapses in adulthood

PNNs are thought to regulate the formation of synapses on the cells they encompass (Ruzicka et al., 2022), and given that the excitatory/inhibitory balance in the PFC is remodeled during adolescence (Caballero et al., 2021), we were interested in assessing whether AIE resulted in alterations of excitatory/inhibitory puncta synapsing on PNNs. Using immunofluorescence and confocal imaging, we quantified the number of vGlut2 and vGat puncta proximal to PNN surfaces (Fig. 2B-D). A 2 (adolescent exposure) x 2 (sex) ANOVA of vGlut2 expressing synaptic puncta identified a significant main effect of exposure, F (1,38) = 7.04, *p* = 0.01, and an interaction of adolescent exposure and sex, F (1,38) = 3.97, *p* = 0.05. As seen in Figure 2E, there was a tendency for AIE-exposed animals to have lower vGlut2 puncta proximal to PNNs, with this effect being driven by AIE-exposed females (*p* < 0.05). A two-way ANOVA of the number of vGat expressing puncta per volume of PNNs revealed a main effect of adolescent exposure that approached significance, F (1,37) = 3.71, *p* = 0.06 (Fig. 2F). When we assessed the ratio of excitatory/inhibitory puncta on PNNs, no effects were observed (Fig. 2G).

### Experiment 3: Immediate effects of AIE on WFA-labeled PNNs

To further evaluate whether AIE-induced changes in PNNs could be evident immediately following the adolescent exposure period, we used WFA labeling of PNNs in the PrL of animals euthanized one hour following the final exposure to water or ethanol on P45 (see Fig. 3A). A 2 (adolescent exposure) x 2 (sex) ANOVA of the number of WFA-labeled cells in the PrL revealed a significant interaction of adolescent exposure and sex, F (1,38) = 8.48, *p* < 0.01. Post-hoc comparisons identified fewer WFA+ cells in the PrL of AIE-exposed males relative to water-exposed males (*p* = 0.0140, Fig. 3C).

### Experiment 4: Immediate and long-lasting effects of AIE on PNN-associated CSPG gene expression

To further assess AIE changes in PNN expression during the exposure period or following a period of forced abstinence, we assessed gene expression of CSPGs (Table 1) in the mPFC (PrL and IL) either following the final exposure on P45 or in adulthood on P70 (Fig. 3B). Separate for each gene 2 (adolescent exposure) x 2 (test age) ANOVAs revealed no significant main effects and interactions for Bcan, Pcan, and Vcan (see Table 2 for values). For Acan gene expression, a significant interaction of adolescent exposure and test age was identified, F (1,39) = 5.92, *p* < 0.05. Follow up comparisons revealed greater Acan gene expression AIE-exposed males compared to water-exposed counterparts immediately following the end of the exposure period (P45), *p* < 0.05, as well as a difference between AIE-exposed P45 and P70 males, *p <* 0.01 (Fig. 3D). For Ncan gene expression, analysis revealed a significant main effect of age, F (1,40) = 12.01, *p* < 0.01, with P45 adolescents having higher Ncan gene expression in the mPFC than P70 adults (Fig. 3E). Given limited differences evident with individual PNN components, we compared gene expression of multiple enzymes that degrade PNNs (MMP9, ADAMTS4/5), with analyses also revealing no differences between exposure conditions or ages (see Table 2).

**Table 2.**
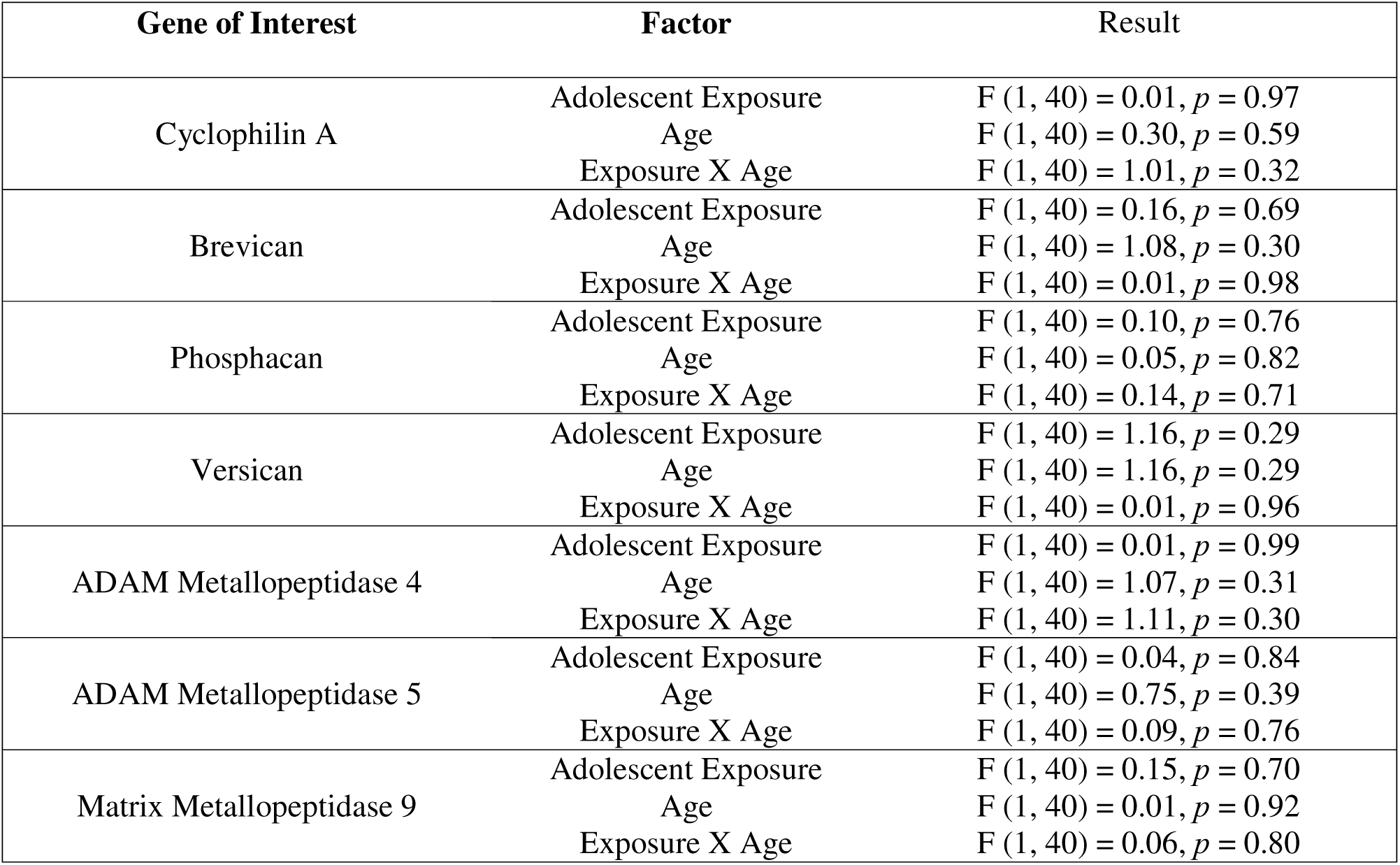
Results of two-way ANOVAs of expression levels for each gene of interest.

### Experiment 5: Validating ChABC-induced PNN depletion and timing of PNN regrowth

ChABC injection leads to a temporary depletion of PNNs, with a gradual restoration of PNN expression evident over time (Lensjø et al., 2017). Our intention was to eliminate the increased number of PNNs in males exposed to AIE and allow for a recovery of PNNs in adulthood (Experiment 6). Therefore, to confirm the efficacy of ChABC and assess the timing of PNN regrowth post-ChABC, we measured WFA labeling at 3 time points after microinjection of ChABC into the PrL (Fig. 4A-C). A one-way ANOVA of the number of WFA+ cells revealed an effect of time post-injection of ChABC, F (2,27) = 49.48, *p* < 0.0001. Follow up comparisons using Bonferroni tests revealed significantly lower WFA labeling in 7-days, compared to both 14- and 21-day post-ChABC animals (*ps* < 0.0001). As seen in Figure 4D, WFA+ cell counts were also lower in the 14-day animals compared to the 21-day subjects (*p* < 0.01).

**Figure 4.**
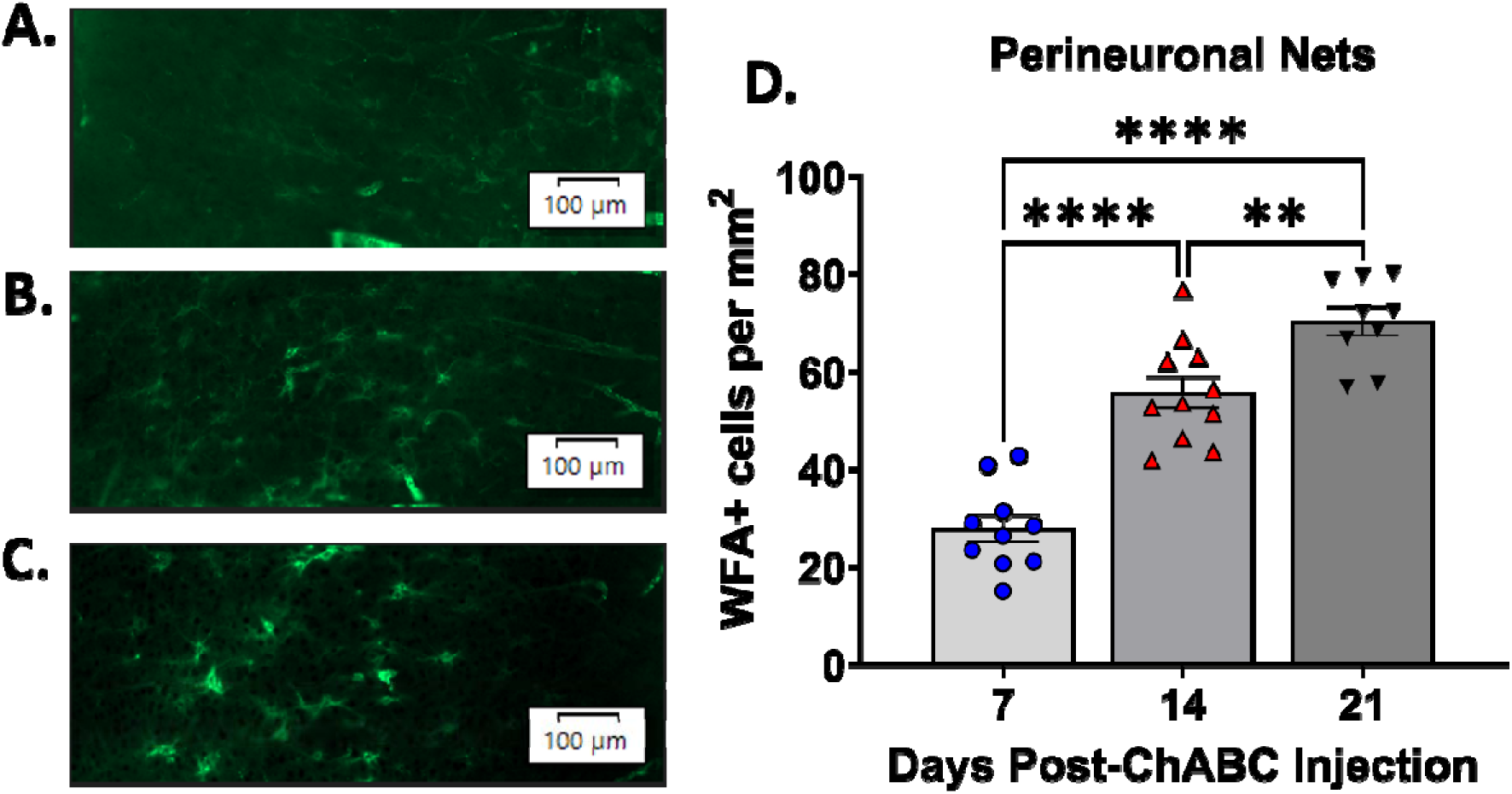
Degradation of PNNs with ChABC and the timing of PNN regrowth. **A-C.** Representative images of PrL WFA labeling in animals one, two, and three weeks following ChABC injection, respectively. **D.** Labeling of PNNs with WFA was the lowest in animals one-week post-ChABC, with gradual increases evident two and three weeks post-ChABC injection. ** *p* < 0.01 and **** *p* < 0.0001 indicate difference between weeks post-ChABC. Scale bar represents 100 μm.

### Experiment 6: Effects of PNN ablation in the PrL on social interaction in adult AIE-exposed rats

Given our finding of increased PNN expression in adult AIE-exposed subjects as well as the role of PNNs in regulation of inhibitory signaling (Carceller et al., 2020), we were interested in determining whether differences in PNN expression contribute to AIE-induced social deficits. To do so, we used the enzyme ChABC to degrade PNNs in the PrL following adolescent exposure and socially tested rats in adulthood (Fig. 5A). A 2 (adolescent exposure) x 2 (drug) ANOVA of male social investigation revealed a significant main effect of drug, F (1,39) = 8.89, *p* < 0.01, with ChABC-induced depletion of PNNs reducing social investigation regardless of adolescent exposure condition (Fig. 5B). Similarly, an ANOVA of the social preference/avoidance coefficient identified a significant main effect of drug, F (1,37) = 5.78, *p* < 0.05, with males injected with ChABC having a lower coefficient, regardless of adolescent exposure (Fig. 5C). Controlling for potential differences in locomotor behavior, a two-way ANOVA of total crosses made during the interaction test revealed a main effect of adolescent exposure, F (1,37) = 7.05, *p* < 0.05, with AIE-exposed males displaying more crosses during the test than their water-exposed counterparts (Fig. 5D). In females, analysis revealed no differences for social investigation, social preference/avoidance coefficient, and locomotor behavior (Fig. 5E-G).

**Figure 5.**
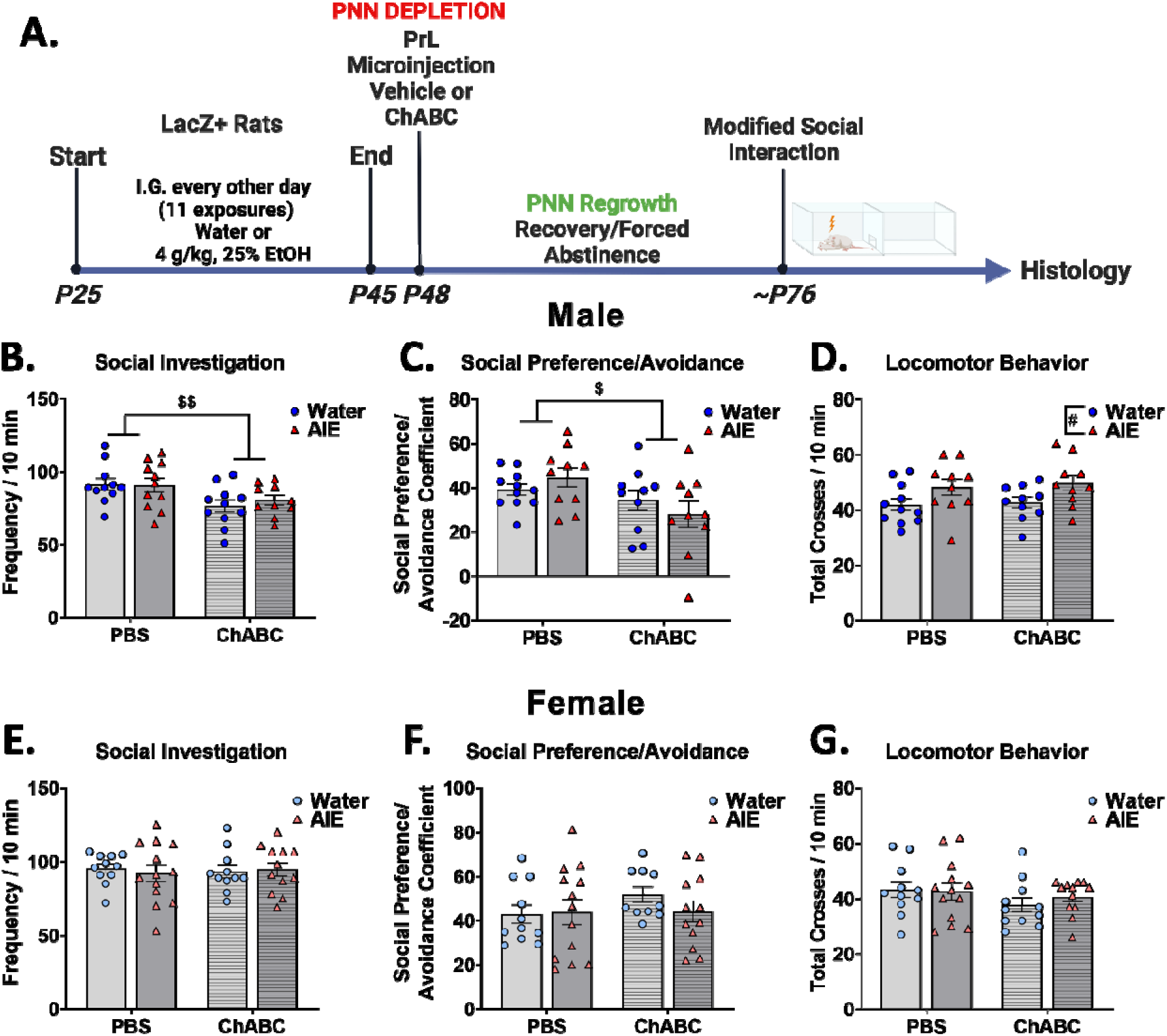
Depletion of PrL PNNs in late adolescent water- and AIE-exposed males and females: Impact on social behavior in adulthood. **A.** PNNs were degraded with ChABC two days after the end of AIE (or water) exposure and allowed to regrow prior to social testing in adulthood. **B-C.** Social investigation and social preference were decreased following ChABC-induced depletion of PNNs in males specifically. **D.** AIE-exposed males displayed greater locomotor behavior, regardless of PNN degradation. **E-G.** Social behavior of females was unaltered following PNN degradation in the PrL. $ *p* < 0.05 and $$ *p* < 0.01 indicate differences between vehicle and ChABC injected subjects.

To evaluate the state of PNNs at the time of behavior, we conducted WFA staining in the PrL and compared groups receiving vehicle or ChABC 28 days prior. A 2 (adolescent exposure) x 2 (drug) ANOVA on WFA labeling in males revealed a significant main effect of drug, F (1,39) = 9.32, *p* < 0.01, with increased WFA+ cells evident in males injected with ChABC, regardless of adolescent exposure (Fig. 6C, representative images Fig. 6A-B). In females, analysis also revealed a significant main effect of drug, F (1,43) = 12.12, *p* = 0.001, with ChABC injected subjects having more WFA+ cells than vehicle-exposed (Fig. 6E).

**Figure 6.**
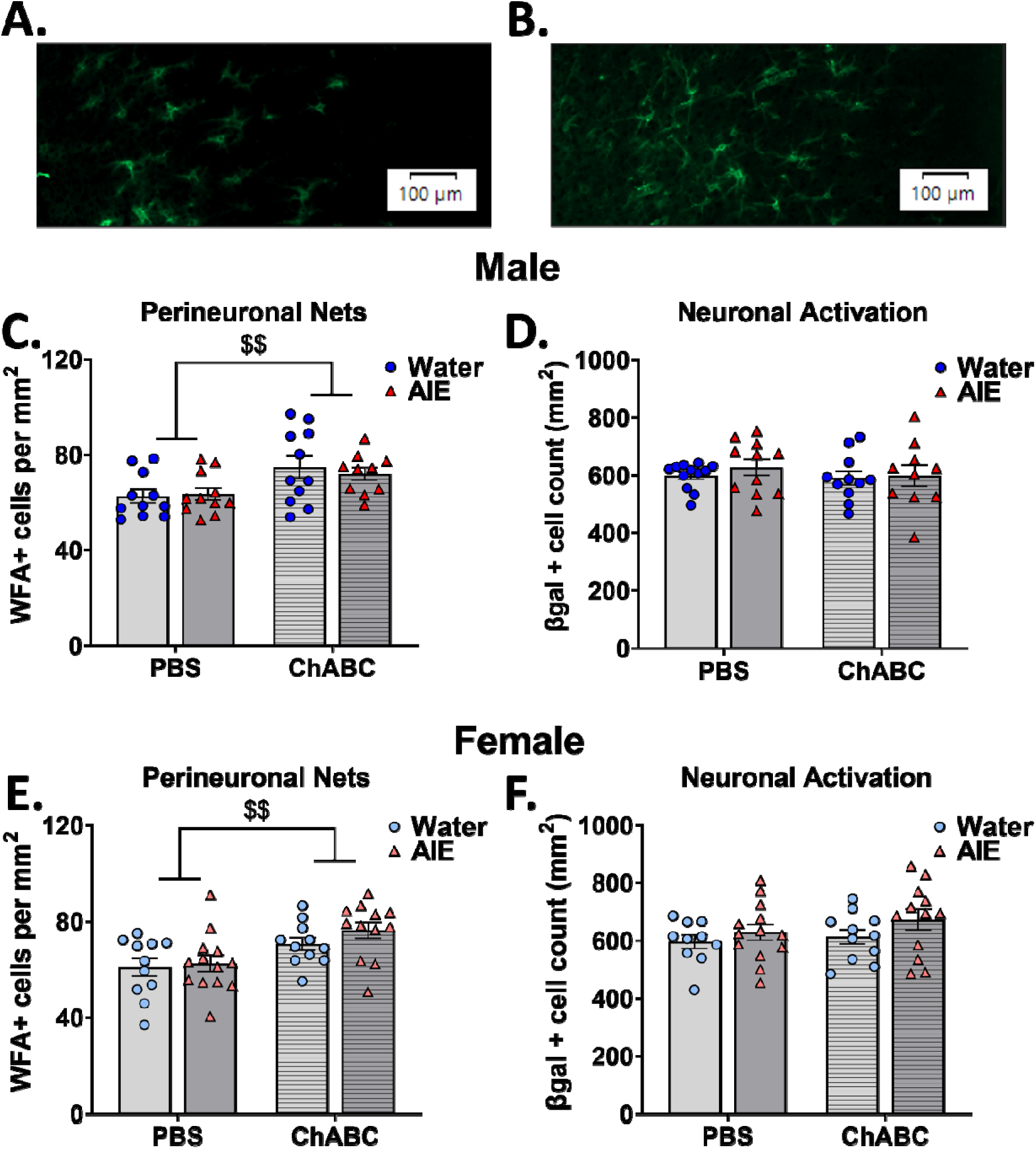
Depletion of PrL PNNs in late adolescent water- and AIE-exposed males and females: Impact on expression of PNNs and neuronal activation of the PrL in adult rats. **A-B.** Representative images of WFA labeling in animals injected with vehicle and ChABC respectively. **C.** Males injected with ChABC 28 days before behavior displayed an increase in WFA labeling in the PrL compared with vehicle injected rats. **D.** No difference in β-gal expression was found between groups. **E.** In females, ChABC injected rats had greater WFA labeling, regardless of adolescent exposure. **F.** Neuronal activation, indicated by β-gal expression, was not affected by PNN degradation in females. $$ *p* < 0.01 indicates difference between vehicle and ChABC injected subjects. Scale bar represents 100 μm.

We also conducted β-gal histology to determine whether PNN ablation and restoration disrupted neuronal activation induced by an interaction with a social stimulus. Two-way ANOVA of β-gal+ cells in the PrL revealed no main effects or interaction for either sex (*p*s > 0.05, Fig. 6D & F).

## Discussion

In the current study, AIE exposure led to a sex-specific increase in PNNs within the PrL of adult males but not females. Additionally, a greater proportion of neurons surrounded by PNNs co-expressed β-gal, suggestive of a greater activation of PV interneurons encapsulated by PNNs in AIE-exposed males that concurrently display social deficits. Further, ablating PNNs during mid/late adolescence revealed a role of PNN alterations in adult social behavior that was not dependent on exposure to water or ethanol during adolescence, thereby supporting a critical role for fine-tuning of PNN expression and PV interneuron function during late adolescence.

Exposure to ethanol during adolescence has been shown to increase the expression of PNNs in the frontal cortex in adulthood (Coleman et al., 2014; Dannenhoffer et al., 2022). Using WFA to label PNNs in the PrL, we similarly found that AIE exposure increased PNN expression, replicating previously reported findings (Dannenhoffer et al., 2022). These findings extend previous reports in that the effect of AIE exposure on PNN expression was sex-specific and evident only in males. Previous studies that found AIE-induced changes of PNNs used only male subjects; therefore, this is the first study to show female subjects are insensitive to AIE-induced changes in PNN expression, at least within the PrL. Future studies will need to determine sex specificity in previously identified regions that are altered by ethanol. Similar sex-specific PNN alterations resulting from juvenile and adolescent manipulations such as unpredictable stress (Page & Coutellier, 2018) and social isolation (Gildawie et al., 2021) have been identified. PNNs rapidly increase during the juvenile/adolescent transition to adulthood, thus it is not surprising that insults such as ethanol and stress during this ontogenetic maturation of PNNs impairs PNN expression levels later in life. However, females have been reported to undergo an earlier ontogenetic increase in PNNs, thought to be associated with earlier pubertal timing in these subjects (Drzewiecki et al., 2020), a finding that may be a contributing factor to the observed sex differences.

The upregulation of PNNs in AIE-exposed males could be the result of several factors. PNNs protect PV interneurons from insults such as oxidative stress (Cabungcal et al., 2013) and inflammation (Egea et al., 2010). AIE influences both oxidative stress (Pelição et al., 2016) and neuroimmune response (Crews et al., 2019) making it possible that increases in PNNs during protracted withdrawal act as a protective measure. Importantly, these effects of AIE exposure on oxidation (Fagundes et al., 2016; Fernandes et al., 2018) and neuroimmune response (Vore et al., 2021) do not appear to be sex-specific, therefore it is unknown why males but not females display PNN upregulation after exposure to AIE. In addition, the potential decrease in PNNs immediately following AIE suggests that PNNs are not immediately increased in response to ethanol directly, therefore more work is needed to identify the protracted withdrawal mechanism mediating this sex-specific dysregulation of PNNs after AIE.

In addition to PNN expression being upregulated after AIE exposure, we also found more PNN encompassed cells to be activated by an interaction with a social partner in AIE-exposed males. Importantly, AIE-exposed males also displayed social deficits, and the greater activation of PNN encompassed PV interneurons in these subjects suggests these specific cells may contribute to social impairments. These findings are supported by previous studies showing that chemogenetic activation of PV interneurons in the PrL led to reduced social preference in adult male rats (Ferguson & Gao, 2018b). Taken together, this suggests a prominent role for PV interneuron functioning in social responses. It is likely that PV interneuron functioning must be finely tuned, as studies in adult male mice have also shown increased PV interneurons firing rates during exploration of a social partner, with chemogenetic inhibition of these interneurons reducing sociability and activation enhancing prosocial responding (Liu et al., 2020). These opposing findings suggest that too much or too little PV interneuron activity may result in social deficits. PV interneurons significantly contribute to the excitatory/inhibitory balance (Ferguson & Gao, 2018a) by regulating pyramidal neuron output; therefore it is likely that the greater activation of these cells in AIE-exposed males dampens pyramidal neuron activity. In fact, previous evidence supports hypoactivity of pyramidal neurons being associated with social deficits in rodent models of autism spectrum disorder (Asgarihafshejani et al., 2019; Brumback et al., 2018; Howell et al., 2017; Sacai et al., 2020).

We previously found differential activation of the PrL between water- and AIE-exposed males (Towner et al., 2023), however, in the current study, β-gal+ cell counts in the PrL were similar in water-exposed and AIE-exposed males. This discrepancy between the two studies is likely due to methodological differences. In our previous study, we focused on neuronal activation of superficial layers II/III of the PrL, however, given greater proportions of PNNs in deeper layers (Alpar et al., 2006; Lipachev et al., 2019), the current study assessed expression of both PNNs and β-gal across all layers. The additional β-gal expression in layers beyond II/III could have eliminated previously observed differences in β-gal+ cell counts between AIE-exposed and water-exposed, suggesting a potential layer specific effect. Further evaluation of layer differences is warranted.

In contrast to males, females exposed to AIE did not display greater colocalization of WFA and β-gal in the PrL. Importantly, female social behavior also was not affected by AIE (Towner et al., 2023). If these activated PV interneurons contribute to social deficits it is therefore not surprising that differences in WFA and β-gal colocalization were absent in females. The role of the PrL in regulating social investigation may be sex-specific, a possibility supported by our previous findings of PrL inactivation reducing social investigation in male rats but not their female counterparts (Towner et al., 2023). Similarly, chemogenetic inhibition of the basolateral amygdala to PrL circuit has been found to reduce social investigation in males but not females (Przybysz et al., 2023).

VGlut2 and vGat synaptic puncta on the surface of PNNs was not altered by AIE-exposure in males, suggesting a limited effect of PNN upregulation on the excitatory/inhibitory inputs to PNN encompassed cells. Alternatively, AIE exposure in females, in the absence of PNN expression differences, led to reduced vGlut2 synaptic puncta, potentially disrupting the shift in excitatory/inhibitory balance occurring during adolescence (Caballero et al., 2021). Previously reported female-specific effects of adolescent ethanol include increased glutamate transmission in the basolateral amygdala (Przybysz et al., 2021) and altered metabotropic glutamate receptor function (Kasten et al., 2020). It is important to consider that our finding with vGlut2 puncta were distinctly expressed on PNNs and may not represent a general change in vGlut2 expression across the PrL. The decrease in vGlut2 in females suggests that PNN encompassed PV interneurons may receive less excitatory signal and therefore are less likely to mediate output of the PrL, potentially making AIE-exposed females prone to other behavioral alterations such as increases in non-social anxiety-like behavior (Sakharkar et al., 2019; Varlinskaya et al., 2020).

The lower PNN expression identified immediately following the final AIE exposure clearly demonstrates that AIE-associated upregulation of PNNs in the PrL was specific to adult rats that underwent 25 days of forced abstinence. Counter to our original prediction, males exposed to AIE had lower WFA+ cell counts, suggestive of an immediate, short-term blunting of PNN development by adolescent ethanol exposure. Alternatively, AIE exposure in male could directly degrage PNNs similar to the effect of juvenile/adolescent stress (Gildawie et al., 2021; Page & Coutellier, 2018) and other drugs of abuse (for review see Lasek et al., 2018). In contrast to the effect of ethanol exposure during adolescence, six weeks of ethanol drinking in adult mice without a period of forced abstinence increased PNNs in the insular cortex, (Chen et al., 2015), suggesting that PNN expression may be directly associated with the timing of ethanol exposure or regional specificity. Taken together, these results suggest that AIE exposure produces differential immediate effects on PrL PNNs than ethanol exposure in adulthood. Furthermore, adolescent stress such as social isolation in mice (Ueno et al., 2017) and footshock in rats (Gomes et al., 2020) leads to a reduction in PNN expression in late adolescence and early adulthood. An identical stressor in adulthood was found to not influence PNN expression (Gomes et al., 2020), further supporting a potential age difference in the immediate effect of stressors, including ethanol. It is also possible that the increased WFA expression in AIE-exposed males after abstinence may be due to delayed development of PNNs and subsequent overcompensation, although more work is needed to confirm this prediction.

AIE exposure led to greater Acan gene expression following the final exposure, however this effect did not persist into adulthood. The upregulation of Acan gene expression and decreased WFA+ PNNs at P45 suggests that Acan gene expression may be compensating for the low levels of PNN protein. However, Acan is only a single component of PNNs and the other CSPG genes assessed were not influenced by AIE. Others have also found no change in CSPG gene expression but altered CSPG protein levels (Van Den Oever et al., 2010). As mentioned prior, abstinence is likely an important contributor to ethanol-induced PNN changes evident in adulthood, and had we assessed CSPG gene expression during abstinence, it is possible that we would have observed differences between exposure groups. It is also worth considering that the tissue was collected from the mPFC, including both the PrL and IL, and exposure to chronic stress in adolescence influences Ncan levels in the PrL but not the IL (Yu et al., 2022), suggesting PNN dysregulation may be subregion specific. It is also important to point out that discrepancy between transcript levels and WFA staining may highlight posttranscriptional effects. We did observe greater Ncan gene expression in adolescent males, supportive of developmental changes in PNN expression as described previously (Baker et al., 2017; Drzewiecki et al., 2020; Yu et al., 2022). Gene expression was only assessed in males, and we cannot conclude that AIE exposure in females elicits similar effects; therefore investigation of potential sex differences is needed.

PrL PV interneurons contribute to social behavior (Ferguson & Gao, 2018b; Liu et al., 2020) and are preferentially encapsulated by PNNs, therefore we predicted that social deficits after AIE would be associated with altered expression PNNs. However, instead of determining if the absence of PNNs would reverse AIE-induced social impairments, we performed social testing after restoration of previously ablated PNNs. If ethanol was no longer present, we predicted that PNNs would return to neurotypical levels and functionally modulate social behavior similarly in both adolescent exposure conditions. Contrary to our prediction, adult males exposed to water and AIE during adolescence displayed reduced social investigation and social preference following PNN degradation and restoration, suggesting that alterations of PNNs in mid/late adolescence impairs adult social behavior. In combination with the effect of AIE exposure resulting in short-term reductions in PNN expression, these data imply that the maturation of PNNs during adolescence is an important process for normative social behavior and that insults (AIE or ChABC) during this developmental period may result in aberrant PNN expression/function and social deficits. In females, altering PNNs had no effect on social behavior.

These findings are the first to provide evidence for a sex-specific role of PrL PNNs in modulating social behavior. A recent study assessing PNN contribution to social deficits in a mouse model of autism spectrum disorder found contrasting results, wherein PNN depletion in the cerebellum ameliorated social deficits (Liu et al., 2023). However, social behavior was observed in the absence of PNNs (Liu et al., 2023), whereas our study allowed for at least a partial restoration of PNNs prior to behavioral testing. Furthermore, as noted prior, adolescence may be critical for the development of proper social behavior, and therefore, manipulations to PNNs in adolescents and adults likely results in different outcomes. It should also be considered that the PrL and the cerebellum as well as PNNs within these regions likely differ in their regulation of social responses.

Surprisingly, both sexes and exposure groups that experienced PNNs depletion with ChABC in mid/late adolescence displayed an increase in PNN expression in adulthood. As was seen in Experiment 1, males that had lower social investigation also had greater PNN expression, further supporting our conclusion that an upregulation of PNNs within the PrL likely contribute to social deficits. Importantly, although PNN expression was also increased after depletion in females, no effects of PNN depletioin on social behavior were observed, suggesting that PNNs within this region play a limited role, if any, in social behavior of females. As far as we know, no prior studies have found ChABC-induced PNN ablation to be associated with a greater expression of PNNs following recovery. Since PNNs were depleted during adolescence, it is likely that the ontogenetic increase in PNNs occurred with overcompensation. In contrast to PNNs, β-gal expression was not altered by ChABC injection, suggesting that the upregulation of PNNs may not have influenced PrL neuronal activation induced by interaction with a social partner.

A few limitations of the current study are worth noting. One of our primary goals was to reverse AIE-induced social deficits with PNN restoration after ablation. However, in our vehicle-treated males, we failed to observe social deficits, thus we cannot conclude whether PNN changes differentially contributed to social behavior in water-versus AIE-exposed males. However, it is more than likely that surgery during mid/late adolescence led to unintended social consequences in water-exposed controls due to adolescent exposure to isoflurane during the procedure (Landin et al., 2019). A second limitation is that the histology of WFA and β-gal after ChABC was done in separate tissue sections, therefore we cannot conclude whether we altered the activation of PNN encompassed cells. A final limitation is that we do not know whether PNNs were restored to equivalent functional levels. Although we observed an increased number of WFA-labeled PNNs in ChABC-injected groups relative to vehicle-injected counterparts, it is possible that these recently developed nets are immature, and we have no evidence as to whether the number and distribution of synapses on PNN encompassed interneurons as well as PV expression were restored.

The current work identified AIE-associated sex-specific alterations of PrL PNN expression. The increase in PNNs was evident in AIE-exposed adult males after a period of abstinence but not immediately after AIE during late adolescence. These alterations were not associated with changes in excitatory or inhibitory synaptic inputs. Temporary degradation of PNNs in late adolescence resulted in social deficits evident in adult males following PNN restoration and subsequent upregulation. Collectively, these findings suggest that AIE dysregulates PNNs in the PrL which contributes to AIE-induced social impairments.

## Acknowledgements

Supported by NIH grants U01 AA019972 (Neurobiology of adolescent drinking in adulthood, NADIA), T32 AA025606 (Development and Neuroadaptation in Alcohol and Addictions, DNAA) and F31 AA029300. Any opinions, findings, and conclusions or recommendations expressed in this material are those of the author(s) and do not necessarily reflect the views of the above-stated funding agencies. The authors have no conflicts of interest to declare.

